# CGBack: Diffusion Model for Backmapping Large-Scale and Complex Coarse-Grained Molecular Systems

**DOI:** 10.1101/2025.06.04.657965

**Authors:** Diego Ugarte La Torre, Yuji Sugita

## Abstract

Molecular dynamics simulations based on coarse-grained (CG) models are used to accelerate conformational dynamics of biomolecules and other chemical systems with reduced computational costs. CG models achieve it by discarding atomic information necessary for downstream structural analysis. Recovering the atomic detail from CG structures, i.e. backmapping, remains a fundamental challenge in multiscale modeling, especially for proteins and complex biomolecular assemblies. Recent machine learning methods have shown promise in reconstructing atomistic details, but most approaches only target simple systems at a small scale. In particular, conventional backmapping pipelines often fail to preserve stereochemistry, induce high-energy configurations, or require extensive minimization. Here we present CGBack, a backmapping framework that employs a denoising diffusion probabilistic model to reconstruct all-atom molecular structures from CG representations. CGBack consists of backmapping and refinement procedures and accurately recovers atomic coordinates across diverse protein systems from small to large scales. We show that CGBack is capable of accurately backmapping both single-chain and multi-chain molecular systems, including densely packed intrinsically disordered proteins in condensates. These results suggest that CGBack can be a powerful tool for multiscale molecular simulation pipelines. We anticipate that CGBack will enable more efficient workflows for protein modelling, as well as for other biomolecules across different CG models.

## INTRODUCTION

All-atom (AA) molecular dynamics (MD) simulations are extensively used across disciplines such as chemistry^1^, biophysics^2^, and materials science^3^ to explore the interactions and motions of diverse molecular systems at an atomic detail. These simulations treat each atom explicitly, accounting for all intra- and intermolecular forces governed by empirical or ab initio force fields. However, the applications of AA models to simulate large systems remain severely limited because of the high computational cost and limited scalability. As target molecular systems increase in size and complexity, the number of atoms involved scales rapidly, often reaching millions or more particles. Simulating such systems with atomic detail not only demands massive computing resources but also constraints the simulation length and the sampling efficiency. To overcome the limitations of AA models, coarse-grained (CG) models have emerged as an alternative to extend the spatiotemporal reach of AA MD.^4, 5^ In CG modeling, groups of atoms are represented as a single interaction site or bead, significantly reducing the degrees of freedom in a system. This simplification enables much larger integration time steps and longer simulation trajectories, facilitating the exploration of large systems that would be otherwise inaccessible using atomistic approaches.^6^

The abstraction of atomic detail in CG models introduces fundamental trade-offs. Many physical interactions, like hydrogen bonding, side chain rotamer states, or solvation effects, are either approximated or entirely absent. As a result, while CG simulations provide valuable mechanistic insights at large scales, they are inherently limited in their capacity to answer questions that depend on atomic specificity.^7^ Bridging this resolution gap, particularly through reversible transformations between CG and AA representations, has become an essential requirement in multiscale simulation workflows. This requirement has led to backmapping,^8^ a process of reconstructing AA structures from CG snapshots or trajectories. In typical multiscale workflows, extensive sampling is first performed using CG models to identify representative conformational ensembles or rare-event transitions. These CG structures are then backmapped into AA resolution to allow detailed structural, energetic, or dynamic analysis through subsequent atomistic simulations, energy minimizations, or machine learning–driven prediction tasks.^9^ In other applications, backmapping is used to initialize MD simulations with realistic large-scale configurations obtained from CG modeling.^10, 11^ Yet, reliable backmapping remains a challenging task. Unlike coarse graining, which follows well-defined mapping schemes, backmapping is inherently underdetermined: multiple AA configurations can correspond to a single CG structure. Therefore, successful backmapping must recover atomic-level detail while maintaining physical plausibility and consistency with the CG input. Inaccurate reconstruction can introduce geometric artifacts, highly energetic configurations, or unrealistic conformations that compromise subsequent analysis or simulations.

Before the advent of deep-learning-based methods, backmapping was primarily addressed using deterministic, rule-based approaches grounded in chemical heuristics, geometric reconstruction, and energy minimization.^8, 12–19^ These tools have served as the backbone of multiscale simulation workflows for over a decade, offering practical means of reintroducing atomic detail to CG configurations. While effective in specific contexts, these methods rely on static assumptions and handcrafted rules, which often limit their applicability across diverse systems and CG representations. One of the most widely used tools in this category is Backward,^8^ a backmapping pipeline developed for the MARTINI^20^ CG force field. Unlike fragment-based methods, Backward uses a geometric-based approach. It reconstructs atomistic structures by projecting CG coordinates using mapping definitions, followed by geometric corrections and a final relaxation stage involving energy minimization and position-restrained MD simulations. Although Backward performs well for standard lipids and proteins, it has limitations when reconstructing extended protein structures. It depends heavily on accurate user-defined mapping files, and may produce strained geometries that require careful relaxation, especially in complex systems. Another notable tool is PULCHRA,^18^ which was originally designed to rebuild protein sidechains from backbone-only representations. PULCHRA uses knowledge-based rotamer libraries and idealized bond geometries to reconstruct heavy atoms, making it useful for CG models that retain only backbone or Cα positions. However, this can lead to unrealistic topologies which are difficult to resolve via standard energy minimization and may require user intervention.

In response to the limitations of these backmapping tools, attention has recently turned to machine learning (ML) models as a promising tool for reconstructing atomistic detail from coarse-grained representations. Unlike traditional approaches that rely on hand-crafted rules or fragment libraries, ML methods aim to learn complex structure-to-structure mappings directly from data, enabling greater flexibility, adaptability, and potentially, higher accuracy. Some of the earliest ML implementations explored diverse neural networks architectures to model the backmapping of CG structures.^21, 22^ More recent methods utilize transformer architectures with attention mechanisms,^23^ enabling the model to capture complex spatial relationships. On the other hand, generative models, including generative adversarial networks,^24^ variational autoencoders,^25–27^ diffusion models,^28–32^ and more recently, flow-matching,^33, 34^ are also gaining traction. These models go beyond point predictions and offer probabilistic sampling frameworks that are particularly valuable in backmapping, where multiple valid atomistic configurations can exist for the same coarse-grained input. Despite these advancements, current ML pipelines for backmapping remain limited in several key respects. Most pipelines are not readily generalizable to diverse molecular environments. Many implementations focus exclusively on reconstructing side chains or secondary structures rather than full molecular reconstructions. Another major limitation in deterministic pipelines is the tendency to treat backmapping as a static, one-shot prediction task. In other words, given a CG input, these pipelines generate a single deterministic output without any iterative refinement or feedback loop.

These limitations motivate the development of CGBack, a backmapping framework that employs a denoising diffusion probabilistic model (DDPM) to produce atomistic structures of complex biological systems. In this work, we present CGBack’s design, including model architecture, training datasets, and the validation strategies employed to assess its performance. The Methods section describes the model implementation, along with the preparation of training data and the evaluation metrics used. In the Results section, we compare the performance of CGBack against three representative backmapping tools: (1) PULCHRA,^18^ (2) cg2all,^23^ and (3) FlowBack,^33^ highlighting significant differences. We further evaluate CGBack’s scalability by testing its performance on proteins of varying sizes and demonstrate its utility through the backmapping of a large protein condensate. Finally, in the Discussion and Conclusions, we assess the current limitations of CGBack, provide practical guidelines for end-users, and outline future directions for enhancing the model’s accuracy, generality, and applicability.

## METHODS

### Data collection and preprocessing

For the training of CGBack, we used the SidechainNet^35^ dataset, an extension of the ProteinNet^36^ dataset that includes high-resolution atomic coordinates, sequences, and structural annotations for protein chains. Specifically, we selected the 100% thinning of the CASP12 subset. In CGBack, each training instance corresponds to a residue, where the objective is to reconstruct the structure of the non-hydrogen atoms from its CG representation and local spatial context. While the original dataset contains a large number of residues, we applied filtering criteria to the training split. To be included, a residue had to meet the following conditions: (1) both its preceding and following residues in the sequence space must be present in the structure, and (2) the target residue itself must contain complete heavy atom coordinates, including sidechains and backbone atoms. The adjacent residues were only required to contain Cα positions, since their role is to provide the necessary context for the backmapping, rather than to serve as reconstruction targets. After filtering, we obtained a final dataset consisting of 21,649,653 residues, each representing a single, valid datapoint.

Each residue was used to build a graph, each containing neighboring residues to provide contextual information (Figure 1a). Specifically, all residues with available Cα coordinates located within a specified distance threshold from the central residue’s Cα atom were included as neighboring nodes. The atomic coordinates were locally centered on the Cα position of the target residue, without the application of any global alignment or rotational normalization. In this study, we experimented with three neighborhood cutoff distances: 6 Å, 8 Å, and 10 Å, allowing us to evaluate the impact of local context size on the model performance.

**Figure 1.**
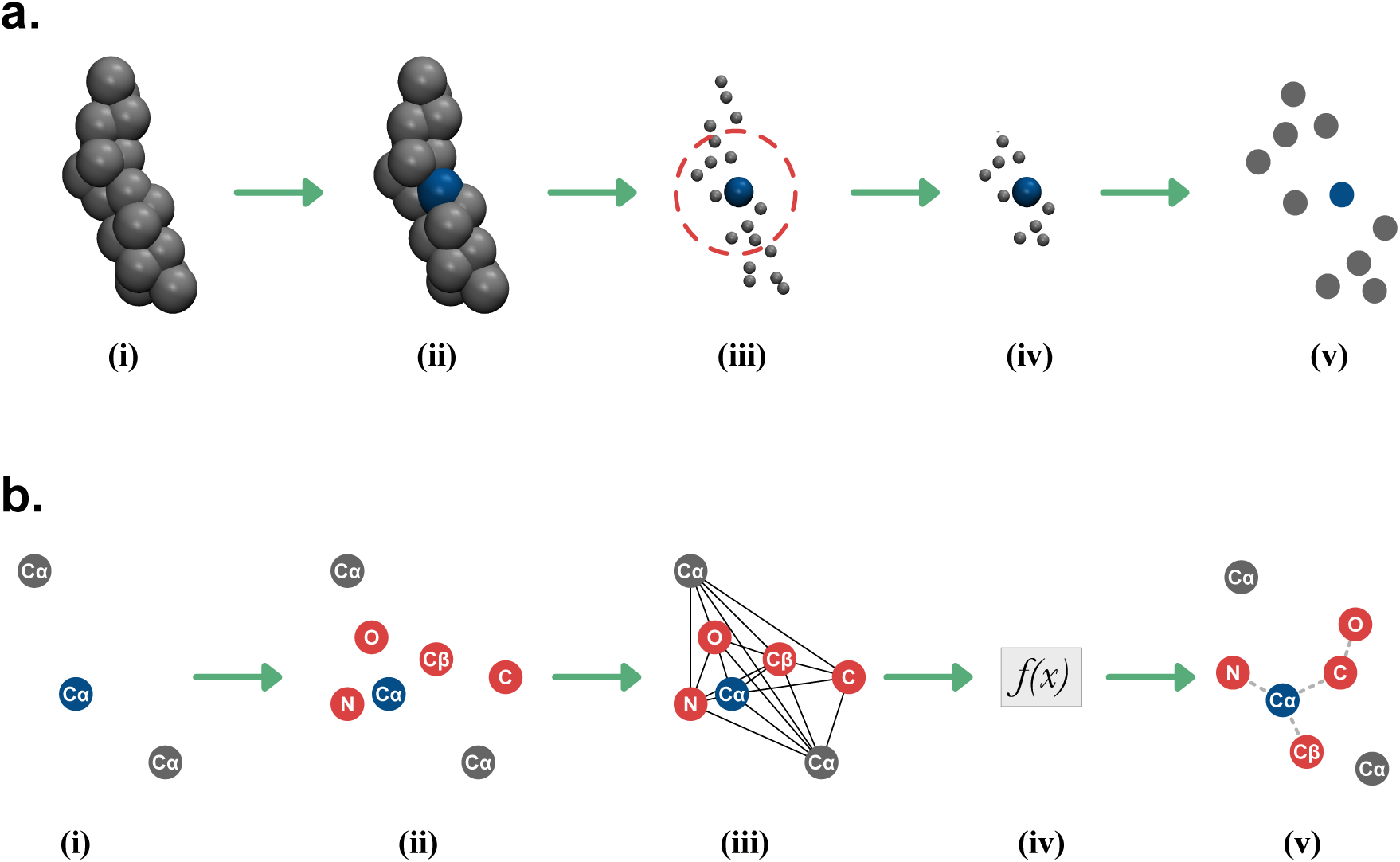
Overview of graph construction and all-atom reconstruction. (a) Graph construction begins with a protein structure represented by Cα atoms (i). A target residue is selected (blue) (ii), and neighboring residues within a defined cutoff distance (indicated by the red dashed circle) are identified (iii). All non-neighboring residues are removed (iv), and the remaining Cα atoms are treated as nodes in a graph (v). (b) All-atom reconstruction proceeds from the constructed graph (i). The missing atoms surrounding the target Cα are randomly initialized (ii), and a fully connected graph is constructed among them (iii). A diffusion model is then applied to denoise these initial positions (iv), yielding a predicted all-atom structure (v). During this process, only the target atoms are updated, while the positions of context atoms (shown in grey) remain fixed. Dashed lines are included for visual guidance only.

To assess CGBack and compare it against existing backmapping methods, we employed the validation and test sets provided in the SidechainNet dataset. Specifically, we used the “valid-90” and “test” splits, which contain 31 and 39 structurally valid proteins, respectively.

Each structure was converted to a CG representation by retaining only the Cα atoms. Both the CG and AA structures were used for performance evaluation across all tested models.

The node features were designed to encode both chemical identity and structural context. Each node captured four components: atom type, residue type, connectivity role, and 3D spatial coordinates. Atom types encompassed all backbone atoms and sidechain atom variants, totaling 36 distinct categories. Residue types were represented across the standard 20 amino acid classes. Connectivity roles encoded the node’s position within the local sequence context: previous, central, next, or unconnected, providing directional cues analogous to a residue index and ensuring the model could distinguish N-to-C terminal orientation. These categorical features were encoded using a one-hot vector. Spatial information was encoded as 3-dimensional coordinates (x, y, z), corresponding to the atom’s real-space position. This resulted in 63 features per node. For the graph structure, all nodes were treated as fully connected, allowing message passing across all pairs without predefined bonding constraints.

### Diffusion model

To reconstruct AA coordinates from CG inputs, CGBack employs a denoising diffusion probabilistic model (DDPM),^37^ which defines a generative process by learning to reverse a progressive noising of atom positions. Let each atomic position be represented as a 3D coordinate 𝒙 ∈ ℝ^3^. The forward process defines a Markov chain that perturbs the data over 𝑇 time steps by gradually adding Gaussian noise:

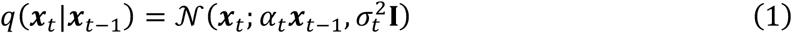

where 𝛼*_t_*𝜖(0,1) and 𝜎*_t_* ∈ (0,1) define a variance-preserving noise schedule, and 𝒙*_t_* is the noisy version of 𝒙 at timestep 𝑡. The goal is to learn the reverse diffusion process:

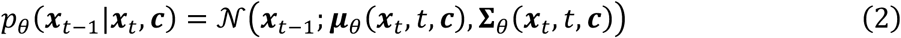

where 𝒄 represents the conditioning information, and 𝝁*_θ_* and 𝚺_θ_ are the parameterized mean and variance, respectively. In practice, the model is trained to predict the noise 𝝐 that was added during the forward process, rather than the denoised data directly. A simplified training objective^37^ is given by:

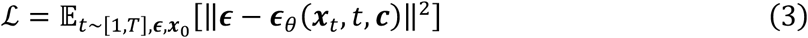

where 𝝐_θ_ is a model that predicts the noise 𝝐, and 𝑡 is sampled from a uniform distribution [1, 𝑇].

At inference time, a sample is generated by drawing 𝒙_𝑇_∼𝒩(0, 𝐈) and iteratively denoising it using the learned network 𝝐_θ_. The denoised coordinate at each step is computed as:

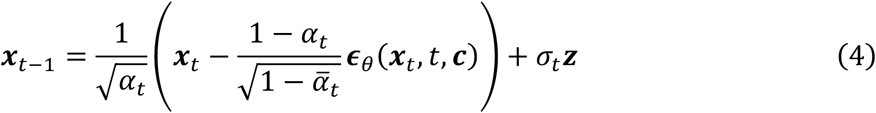

where 𝒛∼𝒩(0, 𝐈) for 𝑡 > 1 and 𝒛 = 𝟎 for 𝑡 = 1, and 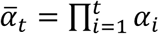. This process continues from 𝑡 = 𝑇 down to 𝑡 = 1, producing the final atomistic configuration consistent with the coarse-grained input (Figure 1b). The denoising is applied only to the target atoms, while the context atoms remain fixed throughout the process.

In this work, we adopt the cosine noise schedule,^38^ which prevents abrupt changes in noise level. The cumulative noise level at time step 𝑡 is defined by:

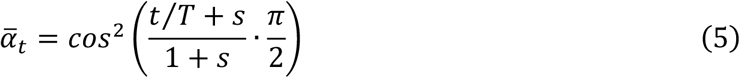

where 𝑇 is the total number of diffusion steps and 𝑠 is a constant set to 0.008. The values of 𝛼*_t_* and 𝜎*_t_* are then derived from 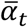 to construct the forward noising schedule.

### Neural network

CGBack employs a graph neural network (GNN) architecture based on complete local frame networks.^39^ Each node 𝑖 in the residue graph is associated with a feature vector 𝒉. ∈ ℝ*^d^* and a coordinate vector 𝒙*_i_* ∈ ℝ^3^, where 𝑑 is the number of scalar features. The node features and coordinates are updated via message passing:

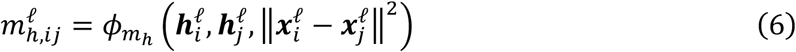

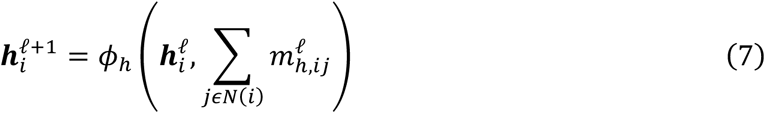

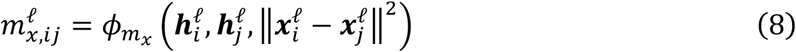

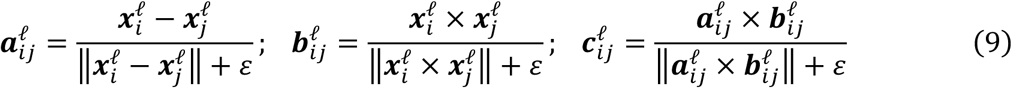

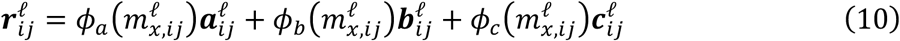

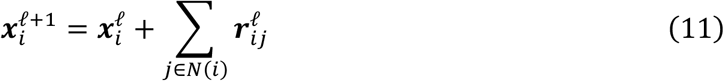

where 𝑁(𝑖) represents the set of neighbors of node 𝑖, ‖ · ‖ is the Euclidean norm, ℓ is the ℓth layer, 𝜀 is a constant set to 1 × 10^-8^, and bold face represent vector quantities. Each function 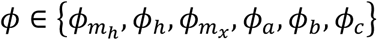 is modeled by a multilayer perceptron (MLP) with the following architecture:

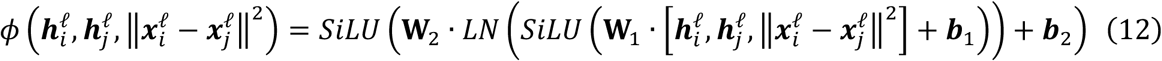

where [ · ] denotes concatenation, 𝐖_1_ ∈ ℝ^d×(2*d*+1)^ and 𝐖_2_ ∈ ℝ*^d^*^×*d*^ are weight matrices, 𝒃_1_, 𝒃_2_ ∈ ℝ*^d^* are bias vectors, 𝐿𝑁 is a layer normalization, and 𝑆𝑖𝐿𝑈 is the Swish function.

### Backmapping pipeline

The CGBack pipeline reconstructs AA structures from CG protein inputs through a three-stage process: (1) backmapping, (2) refinement, and (3) optional energy minimization (Figure 2a). This sequential procedure is designed to recover atomic configurations while correcting structural errors that may arise during the generative process.

**Figure 2.**
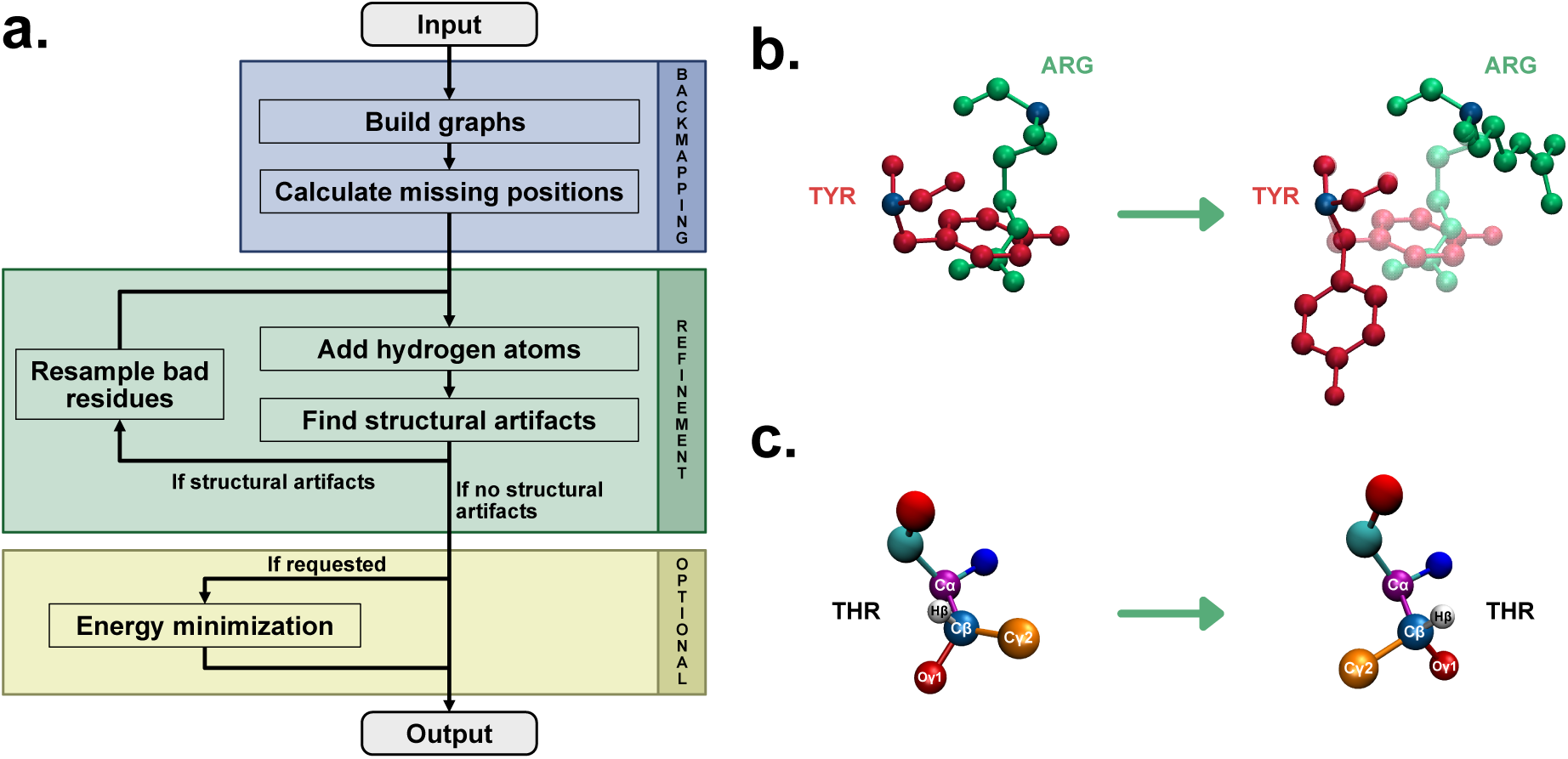
CGBack pipeline. (a) The CGBack workflow consists of three stages: (1) backmapping, (2) refinement, and (3) an optional energy minimization. The refinement stage is executed iteratively until all the structural artifacts are resolved. (b) To resolve ring penetration artifacts, CGBack detects bonds passing through a ring and resamples the atomistic configurations of the residues involved. (c) To resolve chirality errors, CGBack inspects all the stereocenters and resample the residues with an incorrect chirality.

The backmapping stage begins by constructing residue-centered context graphs form CG representations. Each graph is composed of a target residue and its spatial neighbors, selected based on the cutoff distance used during the model training. Then, the heavy atoms corresponding to the atomistic structure are randomly placed following a normal distribution in 3D space. The generative model then iteratively denoises the atom positions by reversing the forward noising process, producing an initial AA reconstruction.

Once the heavy atoms’ final positions are determined, the refinement stage addresses potential geometric and stereochemical anomalies. Hydrogen atoms are first added using simple geometric rules based on atom connectivity. Following hydrogenation, the pipeline perform ring penetration correction (Figure 2b). Ring penetration refers to a steric artifact in which atoms from one residue intrude into the planar region of another’s aromatic or cyclic ring, leading to unrealistic overlaps. CGBack detects these cases by identifying rings within the structure, computing their geometric centroids and normal vectors, and testing whether any bonds from neighboring residues intersect the ring plane. If a bond crosses the plane and its intersection point lies within the projected polygon by the ring atoms, the pair of residues is flagged as penetrated. In such cases, CGBack resamples the involved residues by reapplying the generative diffusion model with varied random noise, effectively generating alternate atomistic configurations until the penetration is resolved. Chirality errors are also detected and corrected during refinement (Figure 2c). Specifically, the Cα atoms of all residues except glycine, and the Cβ atoms of threonine and isoleucine, and determines whether the arrangement of bonded substituents violates the canonical chiral configuration. If an inversion is detected, the affected residue is regenerated using the same resampling strategy to enforce the correct stereochemical configuration. Lastly, the refinement pipeline detects and removes atomic clashes present in the backmapped structure. For users seeking faster processing, this feature can be optionally disabled, especially in workflows where a global energy minimization is performed regardless. This design choice reflects practical considerations, acknowledging that in complex systems, resolving clashes via resampling may take longer than addressing them through routine minimization.

To further improve the structure, CGBack optionally performs energy minimization after refinement. This step relaxes bond lengths, angles, and non-bonded interactions using the L-BFGS^40^ algorithm implemented in OpenMM.^41^ The minimization is performed using the AMBER ff99SB-IDLN^42^ force field combined with the Generalized Born solvent model,^43^ while constraining the positions of the Cα atoms to preserve the original CG backbone. Although optional, this step is beneficial for preparing structures for downstream simulations of modeling tasks.

### Metrics

To evaluate the trained models and compare CGBack’s performance with other models, we employ five metrics: (1) bond score, (2) clash score, (3) ring penetration score, (4) chirality score, and (5) diversity score. Each metric assesses the quality of the generated structures and the generative capabilities of the models.

### Bond score

The bond score measures the quality of covalent bonding within the generated structures. It is defined as the percentage of bonded non-hydrogen atom pairs whose interatomic distance lies within 10% of the bond length specified in the AA reference structures. A score of 100% indicates perfect reconstruction, while lower values reveal increasing occurrences of distorted or unphysical bonds.

### Clash score

The clash score quantifies the prevalence of steric clashes between atoms from different residues. Specifically, it is calculated as the fraction of residues for which one or more of their non-hydrogen atoms fall within 1.2 Å of atoms from another residue. Importantly, for sequentially adjacent residues along the backbone, only sidechain atoms are considered in this calculation to avoid penalizing natural proximity in the backbone. A lower clash score implies physically plausible sidechain packing and spatial arrangements, while a score approaching 100% indicates configurations resembling random atomic placement.

### Ring penetration score

The ring penetration score captures the frequency of geometric artifacts in which bonds intrude into the plane of cyclic groups, typically aromatic rings such as those in tyrosine, phenylalanine, or histidine. These penetrations are detected by computing the centroid and normal vector of each ring and projecting nearby bonds onto this plane. If a bond intersects the plane and its projection lies withing the polygon formed by the ring atoms, it is flagged as a penetration. The score is the computed as the percentage of residues with cyclic groups exhibiting at least one such penetrations. A score of 0% indicates the structure is free from ring penetrations, while higher scores indicate the presence of residues with ring penetrations.

### Chirality score

The chirality score assesses the stereochemical correctness of the reconstructed structures at known chiral centers. For most amino acids, this involves verifying the handedness at the α-carbon (Cα), where the spatial arrangement of the bonded nitrogen (N), carbonyl carbon (C), sidechain β-carbon (Cβ), and α-hydrogen (Hα) must conform to the canonical L-amino acid configuration. The chirality is determined by computing the signed volume formed by these atoms:

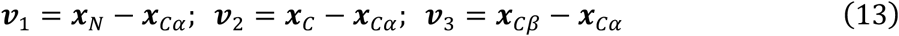

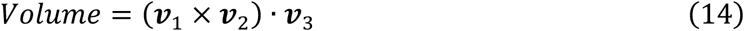

where 𝒙_𝜃_ is the 3D coordinates of atom 𝜃. A positive or zero volume indicates an L-chiral center, while a negative volume corresponds to a D-chiral center. For amino acids with multiple chiral centers, namely isoleucine and threonine, the chirality of the Cβ is also evaluated in a similar fashion. After determining the chirality of all the relevant centers, the chirality score is computed as the percentage of chiral centers that exhibit incorrect stereochemistry. A score of 0% indicates that all chiral centers are correct, while higher values reflect the presence of stereochemical inversions in the structure.

### Diversity score

The diversity score evaluates the generative variability of the model by comparing the structural variance among generated samples to their deviation from a reference conformation.^28, 33^ To compute it, we sample 𝑁 structures from the model and calculate two quantities: (1) the average pairwise root-mean-square deviation (RMSD) among all generated structures (𝑅𝑀𝑆𝐷*_GEN_*), and the average RMSD between each generated structure and the reference structure (𝑅𝑀𝑆𝐷*_REF_*). The diversity score is then given by:

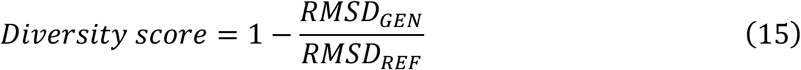

This formulation yields values approximately in the range [0, 1], where 0 indicates maximal diversity and 1 reflects deterministic behavior. In this work we calculate the diversity score using 𝑁 = 10.

### Model training

In this work all neural network components were implemented in Python 3.10.16 using the PyTorch 2.5.0 framework. Data loading was handled using PyTorch’s DataLoader configured with a batch size of 1024, no shuffling, and drop-last enabled to maintain consistent batch sizes. A distributed samples was employed to support multi-GPU parallelization.

Training was conducted on compute nodes equipped with two AMD EPYC 9654 CPUs (2.4 GHz) and four NVIDIA H100 SXM5 GPUs. We used the NVIDIA HPC SDK 25.1, Intel MPI 2021.11, CUDA library 12.8.0, and CUDA driver 570.124.06 in each compute node. All models were trained for 100 epochs using the Adam optimizer with a learning rate of 1 × 10^#H^. To ensure reproducibility, all relevant random number generators (including those from NumPy, PyTorch, and CUDA) were seeded with the value 1705.

After training, each model was evaluated using the checkpoints saved during the final 10 epochs on the validation split. For model selection, we computed a total score based on the following formula:

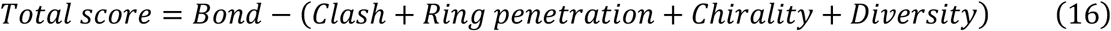

The model achieving the highest total score was selected for downstream analysis and comparative evaluation across models.

### Generation of atomistic structures

To evaluate different trained models of CGBack and to compare CGBack against other backmapping models, we generated atomistic structures for all proteins in the test split of the dataset. For PULCHRA and cg2all, one backmapped structure was generated per protein. In contrast, for FlowBack and CGBack, we generated ten backmapped structures per protein.

For PULCHRA, we used version 3.06 with default settings. For cg2all, we used commit hash 𝑐27𝑑5𝑓0 and the 𝑐𝑜𝑛𝑣𝑒𝑟𝑡_𝑐𝑔2𝑎𝑙𝑙. 𝑝𝑦 script with the options −𝑓𝑖𝑥 and −𝑑𝑒𝑣𝑖𝑐𝑒 𝑐𝑝𝑢. For FlowBack, we used commit hash 𝑐7562𝑓9 and the 𝑒𝑣𝑎𝑙. 𝑝𝑦 script along with the options −𝑣𝑟𝑎𝑚 1, −𝑚𝑎𝑠𝑘_𝑝𝑟𝑖𝑜𝑟, −𝑛_𝑔𝑒𝑛𝑠 10. Additionally, we tested two ODE solvers by setting the −𝑠𝑜𝑙𝑣𝑒𝑟 argument to either 𝑒𝑢𝑙𝑒𝑟 or 𝑒𝑢𝑙𝑒𝑟_𝑐ℎ𝑖. For CGBack, we applied each trained model using its respective hyperparameter configuration to generate the backmapped structures.

### Benchmark

To evaluate the runtime performance of CGBack, we conducted benchmarks on two protein systems with varying size and complexity. The first system is the RC-dLH complex (PDB ID: 7O0W),^44^ and the second is a protein condensate composed of the low-complexity domain of the TAR DNA-binding protein 43 (TDP-43) with amino acid residues 261-414.^45^ For the RC-dLH complex, we generated five subsets of increasing size by sequentially selected the first 1000 to 5000 residues in 1000-residue increments. This setup allowed us to assess how CGBack scales with input size. For each subset, we ran CGBack ten times and report the average runtime and standard error. To assess performance on larger systems, we prepared droplet-like condensates of TDP-43 by computing the center of mass of the structure obtained in previous MD simulations^46^ and selecting chains located within a given radius. We used radii of 300 Å, 400 Å, and 500 Å, corresponding to increasingly complex environments. For each condensate, we ran CGBack ten times and report the average runtime and standard error. Benchmarks were executed on a compute node equipped with two Intel Xeon Gold 6338 (2.0 GHz) CPUs and four NVIDIA A100 GPUs. CGBack was installed with PyTorch with support for CUDA 12.3.0, and the system used CUDA driver version 545.23.08.

### Analysis software

All analyses in this study were conducted using custom Python scripts in combination with the NumPy 2.1.0, SciPy 1.15.0, and Scikit-learn 1.6.1 libraries. Data visualization was performed using Matplotlib 3.10.3, enhanced with the SciencePlots 2.1.1 style package. Molecular visualizations were created using Visual Molecular Dynamics^47^ (VMD) 1.9.4a57 and UCSF ChimeraX^48^ 1.10-rc2025.05.11. All schematic diagrams and illustrative cartoons were designed using Affinity Designer 2.6.3.

## RESULTS

### Hyperparameter tuning

To identify an optimal configuration for CGBack, we systematically evaluated the effect of different hyperparameters on the model performance: (1) the cutoff distance, (2) the number of diffusion steps, (3) the number of GNN layers, and (4) the dimension of the features embedding 𝑑. The effects of these parameters are summarized in Table 1. The cutoff distance determines the spatial neighborhood around each residue. Increasing the cutoff includes more Cα atoms from surrounding residues, enabling the model to better capture non-local interactions and steric constraints. This leads to improvements in the clash and ring penetration scores, decreasing from

**Table 1.** Effect of model hyperparameters on the test split of the CASP12 dataset. Metrics are averaged over 10 generated structures and values are reported as mean ± standard error, except for the diversity score. All the results are calculated before the refinement stage.

| Cutoff<br>(Å) | Steps | Layers | Dimension | Bond<br>(%) | Clash<br>(%) | Ring<br>penetration<br>(%) | Chirality<br>(%) | Diversity |
| --- | --- | --- | --- | --- | --- | --- | --- | --- |
| 6 | 10 | 2 | 128 | 98.94<br>$\pm 0.02$ | 5.27<br>$\pm 0.13$ | 2.32<br>$\pm 0.12$ | 0.048<br>$\pm 0.008$ | 0.071 |
| 8 | | | | 98.76<br>$\pm 0.01$ | 3.21<br>$\pm 0.06$ | 1.42<br>$\pm 0.09$ | 0.100<br>$\pm 0.006$ | 0.064 |
| 10 | | | | 98.57<br>$\pm 0.01$ | 3.27<br>$\pm 0.06$ | 1.61<br>$\pm 0.06$ | 0.126<br>$\pm 0.007$ | 0.036 |
| 8 | 5 | 2 | 128 | 93.82<br>$\pm 0.02$ | 3.01<br>$\pm 0.08$ | 1.33<br>$\pm 0.11$ | 0.063<br>$\pm 0.008$ | 0.096 |
| | 10 | | | 98.76<br>$\pm 0.01$ | 3.21<br>$\pm 0.06$ | 1.42<br>$\pm 0.09$ | 0.100<br>$\pm 0.006$ | 0.064 |
| | 20 | | | 99.58<br>$\pm 0.01$ | 3.55<br>$\pm 0.06$ | 1.42<br>$\pm 0.08$ | 0.062<br>$\pm 0.005$ | 0.026 |
| 8 | 10 | 1 | 128 | 81.29<br>$\pm 0.05$ | 5.74<br>$\pm 0.06$ | 2.29<br>$\pm 0.09$ | 0.732<br>$\pm 0.024$ | 0.069 |
| | | 2 | | 98.76<br>$\pm 0.01$ | 3.21<br>$\pm 0.06$ | 1.42<br>$\pm 0.09$ | 0.100<br>$\pm 0.006$ | 0.064 |
| | | 3 | | 99.50<br>$\pm 0.01$ | 2.56<br>$\pm 0.08$ | 1.00<br>$\pm 0.07$ | 0.021<br>$\pm 0.004$ | 0.016 |
| 8 | 10 | 2 | 64 | 97.81<br>$\pm 0.02$ | 3.54<br>$\pm 0.08$ | 1.58<br>$\pm 0.12$ | 0.118<br>$\pm 0.009$ | 0.072 |
| | | | 128 | 98.76<br>$\pm 0.01$ | 3.21<br>$\pm 0.06$ | 1.42<br>$\pm 0.09$ | 0.100<br>$\pm 0.006$ | 0.064 |
| | | | 256 | 98.92<br>$\pm 0.01$ | 3.07<br>$\pm 0.06$ | 1.19<br>$\pm 0.09$ | 0.068<br>$\pm 0.006$ | 0.032 |

5.27 and 2.32 to 3.27 and 1.61, respectively. The number of diffusion steps governs how gradually the model denoises the atomic coordinates. Longer diffusion processes improve the quality of the reconstructed structures and increase the variability. This is reflected in the increase of the bond and diversity scores. When increasing the number of diffusion steps from 5 to 20, the bond score increases from 93.82 to 99.58 and the diversity score decreases from 0.096 to 0.026. The number of GNN layers controls the model’s capacity to reason over the graph structure. Deeper networks can model more complex spatial dependencies, such as sidechain packing and backbone geometry. Increasing the number of layers from 1 to 2 significantly improved all metrics, and an additional third layer further enhanced the model. Finally, the feature embedding dimension 𝑑 controls the representation of the node features. Starting from an input dimensionality of 63, we observed that increasing the embedding size to 128 substantially improved performance across all metrics. Further increasing the dimension to 256 produced additional, but smaller, improvements, suggesting diminishing returns beyond this point.

Overall, our analyses identified the number of diffusion steps and the number of GNN layers as the two hyperparameters with the most significant influence on the model performance. Based on these findings, we defined three representative model configurations: (1) S, (2) M, and (3) L, corresponding to increasing model complexity and computational cost. All three models share the same cutoff distance of 8 Å, 20 diffusion steps, and an embedding dimension of 128.

They differ in the depth of the GNN: S, M and L use 2, 3 and 4 layers, respectively. The performance of these models is summarized in Table 2. These model configurations were selected to demonstrate the trade-off between computational efficiency and backmapping accuracy. The S model offers the lowest runtime (Figure 3), making it ideal for large-scale or high-throughput applications. The M model strikes a balance between accuracy and efficiency, while the L model achieves the best performance across all metrics, albeit with higher run times. Importantly, the results reported in Table 1 and Table 2 reflect the output immediately after the backmapping stage, prior to the refinement and optional energy minimization steps. In the full CGBack pipeline, refinement removes ring penetrations and corrects chirality errors, while energy minimization resolves any remaining steric clashes.

**Figure 3.**
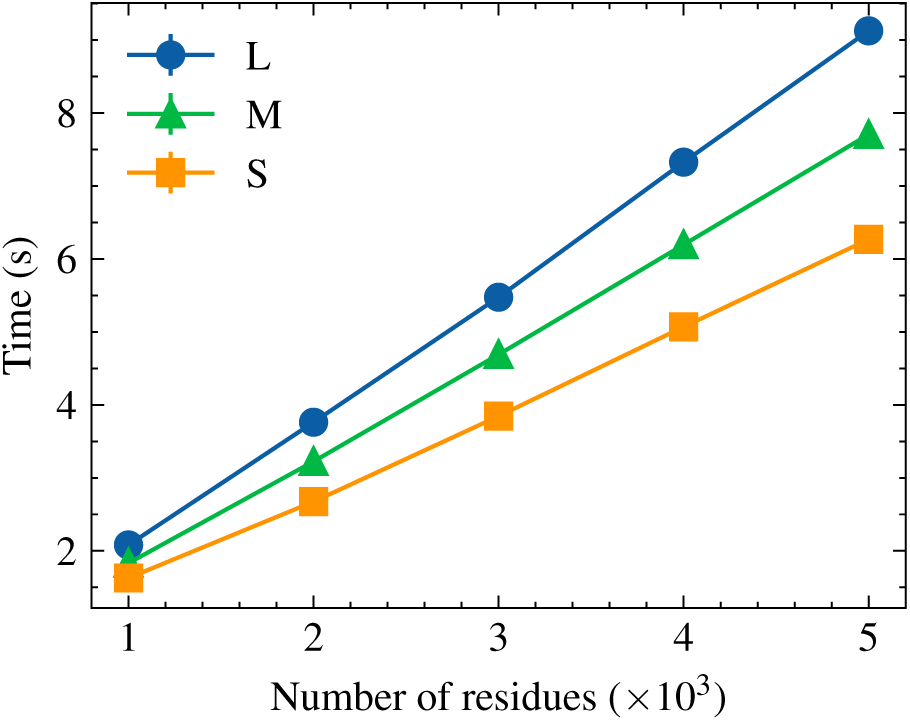
Runtime scaling of CGBack models on the test split of the CASP12 dataset. Backmapping time in seconds against the number of residues for protein structures of increasing size. The error bars represent the standard error. Refinement time is not included.

**Table 2.**
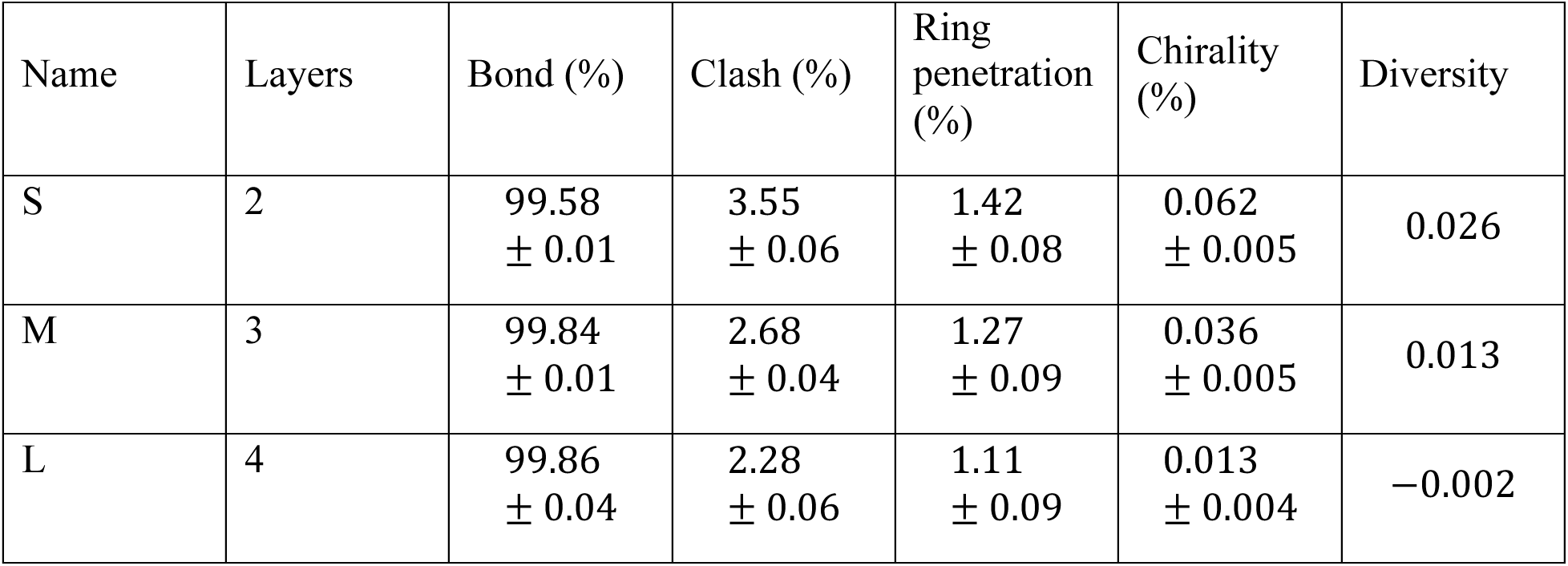
Model performance on the test split of the CASP12 dataset. Metrics are averaged over 10 generated structures and values are reported as mean ± standard error, except for the diversity score. All the models use a cutoff distance of 8 Å, 20 diffusion steps, and an embedding dimension of 128. All the results are calculated before the refinement stage.

### Model comparison

To evaluate CGBack in relation to existing backmapping tools, we compared it with three representative programs: (1) PULCHRA, (2) cg2all, and (3) FlowBack. A summary of performance metrics across all models is presented in Table 3. Both PULCHRA and cg2all demonstrate high bond reconstruction accuracy, with cg2all achieving the best bond score (99.98%), slightly outperforming PULCHRA (98.60%). As a geometry-based heuristic method, PULCHRA produces no chirality artifacts, achieving a score of 0.00%. Similarly, cg2all, which uses an SE(3)-equivariant neural network, preserves the correct stereochemistry. Because PULCHRA and cg2all are deterministic rather than generative models, the diversity metric is not applicable. In terms of ring penetrations, cg2all showed none, while PULCHRA exhibited a small number (0.25%). Although rare, ring penetrations involving non-hydrogen atoms can pose significant issues for downstream energy minimization and simulation stability.

**Table 3.** Comparison of backmapping models on the test split of the CASP12 dataset. For FlowBack and CGBack, scores represent the mean ± standard error. Dash (-) indicates metrics not applicable to deterministic models.

| Model | Bond (%) | Clash (%) | Ring penetration (%) | Chirality (%) | Diversity |
| --- | --- | --- | --- | --- | --- |
| PULCHRA | 98.60 | 0.53 | 0.25 | 0.000 | — |
| cg2all | 99.98 | 0.48 | 0.00 | 0.000 | — |
| FlowBack Euler solver | 99.51 ± 0.01 | 0.09 ± 0.01 | 0.00 ± 0.00 | 3.687 ± 0.050 | 0.219 |
| FlowBack Euler-chi solver | 99.46 ± 0.01 | 0.34 ± 0.02 | 0.04 ± 0.02 | 0.229 ± 0.017 | 0.199 |
| CGBack (M) before refinement | 99.84 ± 0.01 | 2.68 ± 0.04 | 1.27 ± 0.09 | 0.036 ± 0.005 | 0.013 |
| CGBack (M) after refinement | 99.84 ± 0.04 | 2.45 ± 0.03 | 0.00 ± 0.00 | 0.000 ± 0.000 | 0.011 |
| CGBack (M) after full refinement | 99.84 ± 0.04 | 0.00 ± 0.00 | 0.00 ± 0.00 | 0.000 ± 0.000 | 0.013 |

FlowBack represents a generative method based on flow-matching^49^ and an E(3) equivariant GNN (EGNN).^50^ Neural networks built on EGGNs are not equivariant to reflections, causing the model to generate both enantiomers during the structure generation. As a result, FlowBack’s Euler solver yields a high chirality error of 3.687%. Despite this, it achieves a good bond score and has the lowest clash score among all the models (0.09%). To address chirality issues, FlowBack implements an alternative solver, Euler-chi, which applies a correction mechanism during the generative process. This significantly reduces chirality errors to 0.229%, though the fix is only applied to Cα centers and not to Cβ centers, which remained uncorrected. Notably, both FlowBack variants produce structures with almost no ring penetrations, 0.00% and 0.04%, respectively.

For comparison with these models, we used CGBack’s model M, which offers a balance between accuracy and computational efficiency. Before refinement, CGBack achieved a bond score of 99.84%, comparable to FlowBack and PULCHRA, though slightly below cg2all. While its clash and ring penetration scores were initially higher than the other models, refinement with atomic clashes removal turned off eliminated ring penetrations entirely (0.00%). Despite using an SE(3)-equivariant neural network, CGBack still exhibited minor chirality errors (0.036%). In earlier experiments, we tested EGNN-based versions of CGBack but observed a higher frequency of chirality artifacts, which significantly increased refinement time. Switching to a ClofNet-based architecture nearly eliminated these chirality errors. It is worth nothing that although our model is SE(3)-equivariant, the diffusion process itself is not. Recent work on equivariant diffusion models^51^ would help to preserve equivariance throughout the diffusion process and could potentially remove the remaining chirality artifacts. Nevertheless, these chirality errors are fully corrected after applying the refinement pipeline. Additionally, when the full refinement procedure is enabled, which includes clash detection, CGBack eliminates all the structural problems, albeit at the cost of increased refinement cycles. Finally, CGBack exhibits a relatively low diversity score. This facilitates refinement since newly sampled structures are less likely to resemble previously problematic ones, increasing the likelihood of successful correction.

### Large-scale system backmapping

To evaluate CGBack’s scalability and practical applicability to biologically relevant mesoscale systems, we applied it to reconstruct large CG condensates made of TDP-43 proteins (Figure 4a), a system used to model biomolecular liquid-liquid phase separation (LLPS). We generated three condensate systems with varying radii ranging from 300 Å to 500 Å (Figure 4b) and evaluated the quality of the backmapped structures using ring penetration and chirality scores. As shown in Figure 4c, both metrics remained stable and low across increasing system sizes. These results highlight the importance of CGBack’s refinement pipeline, which is particularly effective in correcting steric and stereochemical artifacts even in highly dense and complex systems. Notably, while other models did not exhibit ring penetration issues in the context of single-protein backmapping, such artifacts become apparent in more complex assemblies, including multichain systems. In addition to structural quality, we evaluated the runtime performance. As illustrated in Figure 4d, sampling time scaled linearly with system size, demonstrating strong scalability. The total runtime, including both backmapping and refinement, also followed a linear trend, supporting CGBack’s suitability for large-scale modeling tasks.

**Figure 4.**
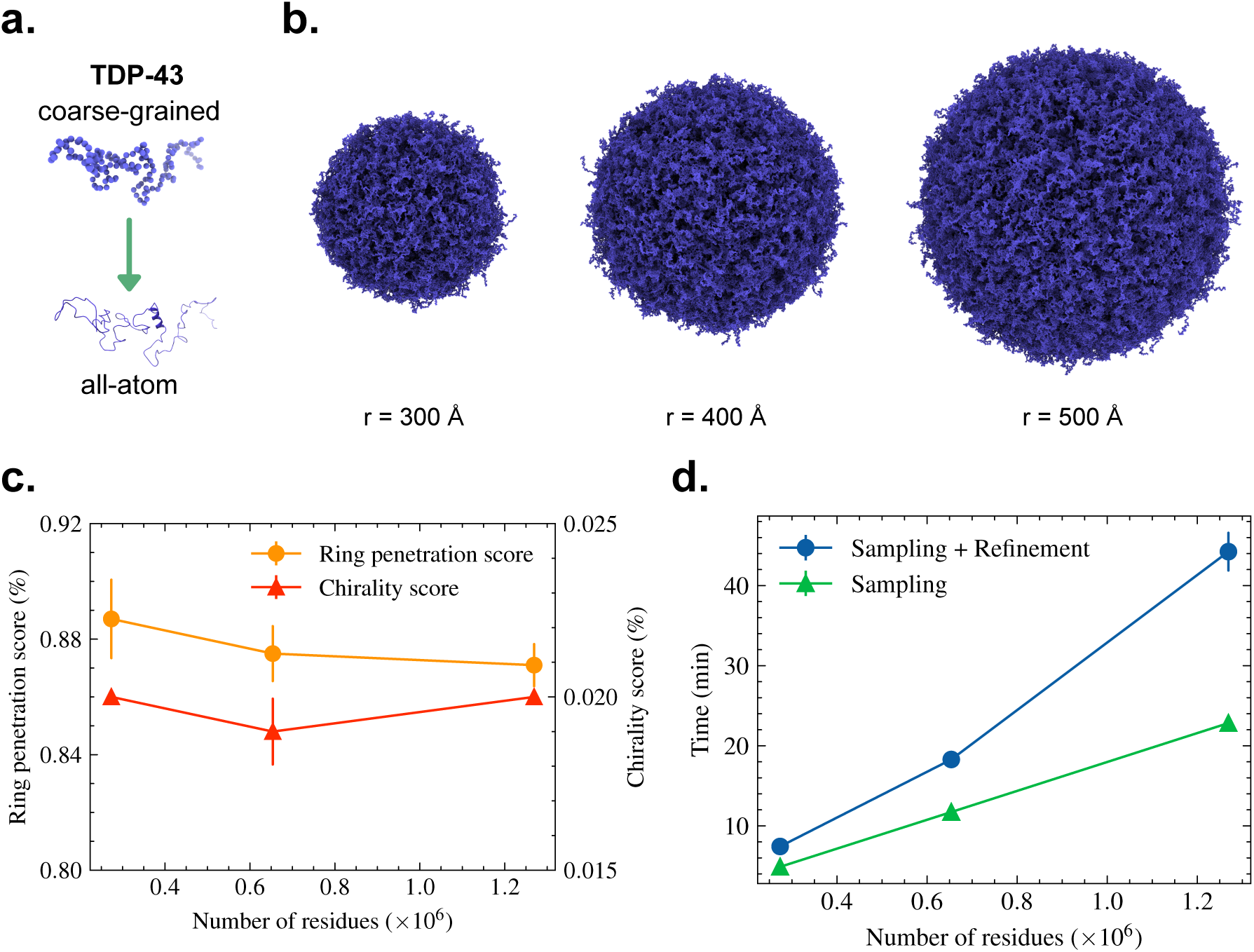
Backmapping of TDP-43 droplet systems. (a) Schematic of both CG and AA representations of a TDP-43 protein. (b) Backmapped TDP-43 condensates of increasing radii. (c) Ring penetration (yellow) and chirality (orange) scores for the backmapped condensates before the refinement process. (d) Benchmark for the backmapping of each condensate. The green line shows the time required for backmapping and the blue line shows the combined time required for backmapping and refinement with atomic clash removal turned off. The error bars in (c) and (d) represent the standard error.

## DISCUSSION AND CONCLUSIONS

In this work, we introduced CGBack, a ML framework for backmapping CG biomolecular structures to their corresponding AA representations. CGBack combines a DDPM with a GNN based on ClofNets to reconstruct atomistic detail from reduced representations. The model exhibits strong performance across diverse scenarios, including both individual protein chains and dense biomolecular condensates. Compared with other tools such as PULCHRA, cg2all, and FlowBack, CGBack offers competitive results across different metrics, including bond score, clash score, ring penetration score, chirality score, and diversity score. A unique feature of CGBack is its integrated refinement pipeline, which systematically corrects geometric and stereochemical artifacts. Additionally, one key strength of CGBack lies in its scalability. It successfully handles large biomolecular systems comprising tens of thousands of atoms and offers flexible model configurations to accommodate both high-throughput and high-accuracy use cases. This adaptability makes it well-suited for integrating it into multiscale simulation pipelines.

Nonetheless, several limitations remain. Currently, CGBack is equipped only with models for backmapping protein systems. Future work will extend its applicability to other biomolecules, including lipids, nucleic acids, and glycans, as well as different coarse-grained resolution levels, such as in the Martini^20^, SPICA,^52, 53^ and iSoLF^54, 55^ force fields for lipids. Additionally, incorporating SE(3)-equivariant denoising could further eliminate stereochemical inconsistencies and move the model closer to generating simulation-ready structures directly from sampling. Another important point of improvement is reducing computation while maintaining structural quality. Since one of CGBack’s objectives is its seamless integration into multiscale MD workflows, enhancing inference speed is a critical goal. Future developments will explore alternative noise schedulers, faster diffusion schemes, and optimized network architectures to enable accurate backmapping with fewer diffusion steps.

Overall, CGBack provides a powerful, generalizable, and efficient solution for backmapping in multiscale molecular simulations. Its ability to recover AA structures from CG representations makes it a valuable addition to the growing toolkit for biomolecular modeling, enabling seamless transitions between simulation resolutions and facilitating downstream atomistic analysis at unprecedented scales.

## DATA AVAILABILITY

The code developed in this study is available at https://github.com/genesis-release-r-ccs/cgback.

The scripts to generate the datasets, the training scripts, and the analysis scripts, as well as instructions on how to use them are available at https://github.com/RikenSugitaLab/cgback-paper. If there is any problem in reproducing our work, please feel free to contact us.

## AUTHOR INFORMATION

### Author Contributions

Diego Ugarte La Torre: Conceptualization (lead); Data curation (lead); Formal analysis (lead); Investigation (lead); Methodology (lead); Software (lead); Validation (lead); Visualization (lead); Writing – original draft (lead); Writing – review & editing (supporting). Yuji Sugita: Conceptualization (supporting); Methodology (supporting); Project administration (lead); Resources (lead); Supervision (lead); Writing – review & editing (lead).

### Notes

The authors declare no competing financial interest.

## ACKNOWLEDGMENT

This research was supported by RIKEN R-CCS and MEXT JSPS KAKENHI [Grant Nos. 21H05249 (to Y.S.)], RIKEN pioneering project “Biology of Intracellular Environments” (to Y.S.), and MEXT program for Big-data-driven bio/synthetic polymer science to create absolutely circular materials (JPMXP1122714694) and Data-Driven Research Methods Development and Materials Innovation Led by Computational Materials Science (JPMXP1020230327) (to Y.S.).

We used computer systems provided by RIKEN R-CCS and the supercomputer Tsubame (hp250065) to train and test our models. We are grateful to Dr. Cheng Tan for sharing the simulation files of the TDP-43 system with us and providing an algorithm for detecting ring penetrations, as well as testing CGBack together with Dr. Yangyang Zhang.

